# AutoNeuro: An Open-Source fMRI Toolbox for Real-Time Neuroadaptive Task Design

**DOI:** 10.64898/2026.04.21.719824

**Authors:** Oliver Sherwood, David Haydock, Raha Razin, Fred Dick, Robert Leech

## Abstract

Real-time functional magnetic resonance imaging (fMRI) offers a powerful means of studying brain function adaptively, enabling experimental parameters to be updated dynamically in response to ongoing neural activity. However, current approaches remain limited by complex infrastructure requirements, bespoke implementations, and a lack of flexible frameworks for closed-loop neuroimaging, with many primarily focusing on neurofeedback experimental designs. Here we present *AutoNeuro*, an open-source framework for real-time fMRI acquisition, preprocessing, feature extraction, and adaptive experimental control. AutoNeuro connects directly to the MRI scanner, receiving reconstructed slices as soon as they become available, and streams them into a modular analysis pipeline designed for low-latency processing. Neural features are estimated at the temporal resolution of acquisition and are passed to a Bayesian optimisation agent that selects task conditions to maximise a user-defined objective function.

Experimental conditions are represented within a bounded “experiment space”, allowing heterogeneous conditions to be explored within a common coordinate system. We demonstrate AutoNeuro in a real-time fMRI experiment in which the system adaptively sampled task conditions to obtain a continuous map of brain response to the range of conditions contained within the experimental space. The system operated within the temporal constraints of real-time preprocessing and analysis, maintaining stable model estimates across iterations, converging on experimental conditions most relevant to the measured brain metric. These results establish AutoNeuro as a flexible platform for closed-loop neuroimaging, supporting hypothesis-driven optimisation as well as exploratory mapping of brain metrics across large experimental spaces.

## 1 Introduction

Most functional Magnetic Resonance Imaging (fMRI) experiments use hypothesis-driven designs with a small set of pre-defined conditions, crafted to isolate specific cognitive processes or brain networks [Huettel et al., 2009]. While highly productive, this framework can obscure the many-to-many mapping between cognitive functions and neural responses [Poldrack, 2006].

In recent years there has been a growing emphasis on alternative, data-driven approaches to mapping functional organisation. Meta-analytic methods [Yarkoni et al., 2011] and machine learning techniques [Varoquaux and Thirion, 2014] have enabled researchers to aggregate evidence across studies and identify broad functional domains. However, these approaches are inherently retrospective: they operate on existing datasets and cannot guide experimental design during data acquisition [Lorenz et al., 2017]. Critically, they enforce a separation between acquisition and analysis. This permits analytical flexibility after data inspection, increasing the risk of post-hoc hypothesis refinement biases.

A different approach is offered by real-time fMRI when combined with neuroadaptive design [Lorenz et al., 2017]. Neuroadaptive design entails analysing neural data as they are acquired and adjusting experimental parameters dynamically based on that measured brain signal. Closed-loop neuroadaptive paradigms eliminate the traditional boundary between acquisition and analysis [Lorenz et al., 2016, 2021].

The most common application of real-time analysis is neurofeedback, a method that uses real-time brain activity to reinforce targeted neural or cognitive states in the participant. Multiple open-access toolboxes exist for neurofeedback applications, or basic real-time frameworks (Supplementary Table 1).

These platforms have substantially advanced the field by providing robust solutions for real-time data streaming, motion correction, ROI-based feedback, and multivariate pattern analysis. For example, Pyneal [MacInnes et al., 2020] and RTPSpy [Misaki et al., 2022] offer accessible Python-based pipelines for real-time preprocessing and feedback control, while OpenNFT [Koush et al., 2017] and AFNI Real-Time [Cox, 1996] provide mature environments with established analysis workflows. More flexible systems such as RT-Cloud [Wallace et al., 2022] enable custom, cloud-based processing pipelines, facilitating distributed and reproducible computation. The FRIEND engine framework [Basilio et al., 2015] offers a well developed neurofeedback suite. Collectively, these tools have lowered the barrier to implementing real-time fMRI and have supported a wide range of neurofeedback and task-based applications.

However, most existing frameworks are optimised for predefined feedback targets or fixed analysis pipelines, rather than for adaptive exploration of experimental parameter spaces. As a result, while they provide powerful infrastructure for real-time processing, they do not natively enable closed-loop neuroadaptive experimental designs.

To this end, Lorenz et al. introduced the “Automatic Neuroscientist” framework, demonstrating how a Bayesian agent can guide task selection based on ongoing neural responses [Lorenz et al., 2016]. This approach, referred to as “neuroadaptive” design, distinguishes itself from neurofeedback by having neural responses guide task presentation, rather than reinforcing some target state in the participant. The approach efficiently identifies task conditions that elicit targeted neural patterns. The approach has revealed functional distinctions such as those between dorsal and ventral frontoparietal networks [Lorenz et al., 2018].

The neuroadaptive approach remains inaccessible to most researchers. Existing implementations are either tailored to specific neurofeedback applications [Basilio et al., 2015], technically complex and bespoke research systems [Goebel, 2012], or focused on isolated demonstrations rather than general-purpose adaptive experimentation. Whilst there has been development of open-access neuroadaptive design frameworks in 2-photon calcium-imaging [Draelos et al., 2025], no open-source platform currently integrates the full real-time pipeline for fMRI— from scanner connection and image reconstruction through preprocessing and feature extraction to adaptive task control—within a flexible and extensible framework.

Here, we introduce *AutoNeuro*, an open-source framework designed to make closed-loop fMRI experimentation widely accessible (Supplementary Table 1, bottom row). AutoNeuro integrates real-time data acquisition, preprocessing, feature extraction, and Bayesian optimisation into a modular pipeline that can be customised for diverse scientific goals. By allowing neural responses to directly guide experimental design, AutoNeuro assists researchers to conceptualise, develop, and run new classes of adaptive experiments, providing a tool to move beyond static task paradigms toward more dynamic, data-driven exploration of multidimensional hypothesis spaces.

We demonstrate AutoNeuro’s capabilities using an experiment defined over a low-dimensional experiment space comprising graded variants of a visual checkerboard paradigm and a motor finger-tapping task. These tasks were selected to elicit robust and dissociable activation in primary visual cortex and primary motor cortex, respectively.

## 2 Materials and Methods

The following section is separated into three parts. First, we describe the architecture of AutoNeuro, explaining the individual modules that allow for a closed-loop, neuroadaptive system to operate (Section 2.1). Second, we describe the agent that is provided as default in AutoNeuro, whilst highlighting that the prospective user can change the agent modularly (Section 2.2). Third, we provide a simple demonstration of AutoNeuro, allowing us to walk through the functionality of the different modules (Section 2.3).

### 2.1 AutoNeuro System Architecture

The AutoNeuro framework is openly available on GitHub, (see hyperlink here), where it is distributed with a comprehensive set-up guide, configuration templates, and a detailed wiki describing system components and usage examples. The framework is designed using existing Python API along with an optional addition of the FreeSurfer algorithm, *SynthSeg* [Billot et al., 2023]. In this section, we will provide an overview of the real-time neuroadaptive system implemented in AutoNeuro, as well as describe the modules that make up the overall design (for t he implementation of the system in a demonstration experiment, see Section 2.3).

AutoNeuro supports an offline acquisition mode for functional neuroimaging data, primarily intended for testing and development. In offline mode, prerecorded fMRI data are streamed volume-by-volume from stored NIfTI files to emulate real-time acquisition, preserving the timing of the original scan sequence by matching the TR in the streaming interval.

#### 2.1.1 System Overview

The neuroadaptive approach to real-time analysis is fundamentally the selection of task or stimulus presentation, based on a metric output from online brain imaging. In order to change the presented task or stimulus to the participant, the researcher must limit the dimensions along which a given task can vary, so as to not make the problem untenable given limited resources and participant constraints. To address this problem, we define an “experiment space” (Section 2.1.3) as the bounded dimensions of task and stimulus variability that AutoNeuro will have access to during scanning (Figure 1, left). Regardless of whether dimensions of the experiment space characterise continuous variance of stimulus (e.g., the frequency of a heard tone), or discrete instances of task conditions (e.g., one-back, two-back), we can characterise the experiment space as a coordinate system, with coordinates representing cross-sections of those dimensions of variability (i.e., 600Hz tone one-back, 2kHz two-back). Across the experiment space, we would expect different landscapes of brain activity, depending on what bounds have been placed on the space, and what metric was taken to summarise brain activity.

**Figure 1:**
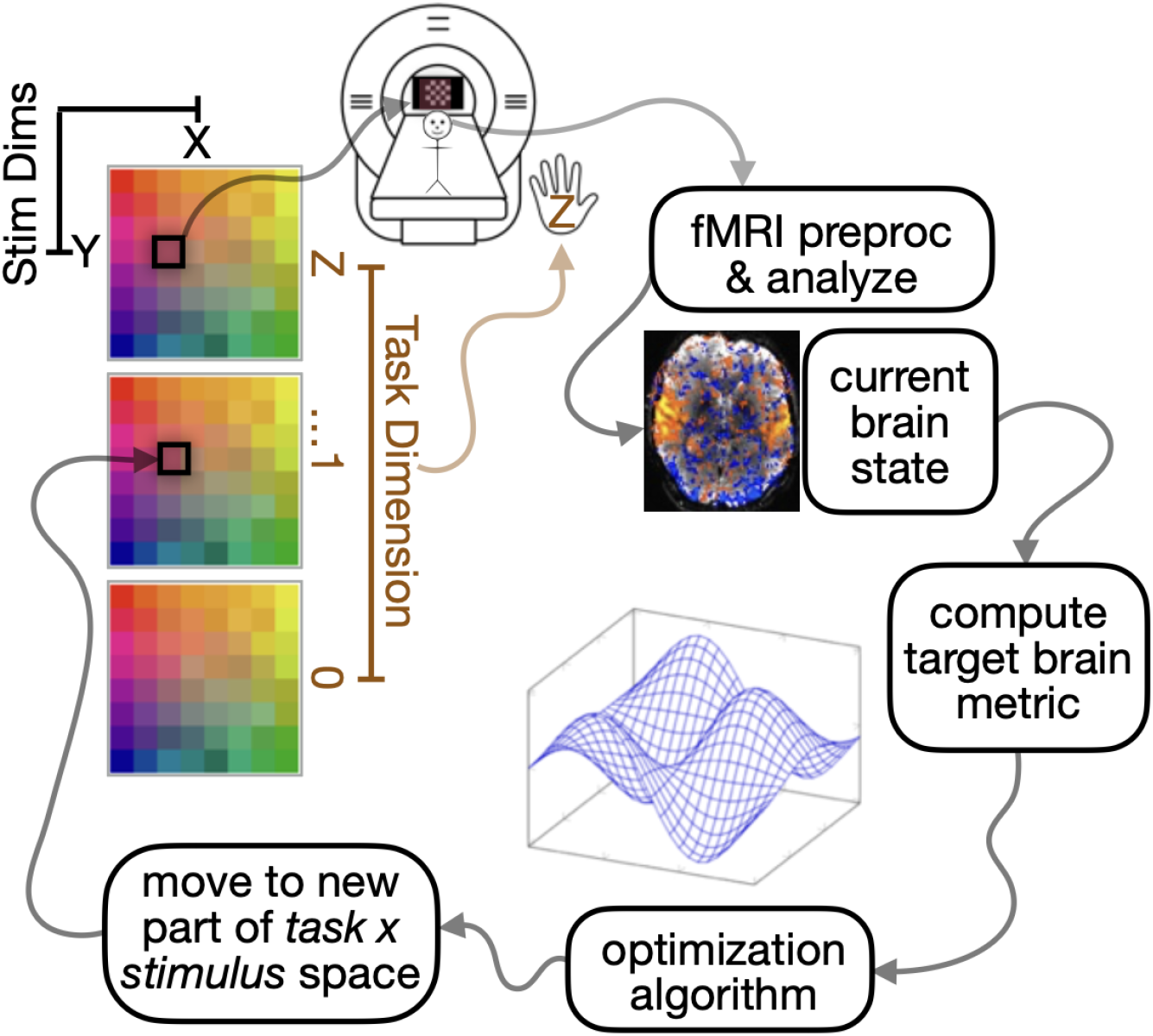
Schematic overview of the closed loop system architecture. Following the process around from the top left, the task and stimulus is chosen from a library of potential experimental contexts in the shared experiment space (represented as coloured matrices, see Section 2.1.3). The participant is presented with an experimental condition from the space. During acquisition, data is preprocessed volume-by-volume, until the end of the experimental block. A target brain metric is computed to summarise the block, which is used in an optimisation algorithm. The output of the algorithm is used to choose the next task/stimulus to present to the participant. As this closed loop iterates, the optimisation algorithm minimises uncertainty about the relationship between the brain metric and the experiment space.

To characterise the space, a participant is presented with a task/stimulus “block” during scanning (Figure 1, top), AutoNeuro takes in the recorded volumes, preprocesses them, and extracts a user-defined feature from the Blood Oxygenation Level Dependent (BOLD) signal, which summarises the neural response during the given experimental block (Figure 1, right).

This metric value is then fed to the “agent” of the system (Figure 1, bottom).

The agent is a model that holds information about the relationship between the experiment space and the brain metric, and uses that information to select the next task to sample in the space. Initially, a small number of user-defined blocks are recorded to train the model, referred to as the “burn-in phase”. This builds a model of the relationship between the experiment space and the target brain metric in the agent. The fit of this internal model is used in the acquisition function (Section 2.3.6) to dynamically suggest the next task and/or stimulus to select from the experiment space in real time.

This completes the closed loop nature of the system which allows the task presentation to be driven by neural response, making the system neuroadaptive. The agent is configured to initially explore the experiment space, sampling points in the space that it is most uncertain about. As the experiment continues, the agent moves from an exploration driven by uncertainty, to an exploitative behaviour, where it targets identification of the point in the experiment space that maximises the brain metric. An intuitive analogy is that of a geological surveyor. Initially, the surveyor samples broadly across terrain to reduce uncertainty about the distribution of resources. As information accumulates, sampling becomes increasingly targeted, focusing on regions predicted to yield the highest density of the target mineral. Similarly, our agent transitions from uncertainty-driven exploration of the experiment space to targeted exploitation of points in the space predicted to maximise the neural response metric.

We note that this is just one application of the neuroadaptive approach. Even the behaviour of explore-exploit is a design choice in this context. The chosen agent behaviour is the means by which the relationship between experiment and brain metric is explored, and can be configured by the user.

#### 2.1.2 System Design

We highlight that the following provided overview of the system design is the default implementation of AutoNeuro, but it is a modular system that can be adapted to accommodate specific user requirements.

The three processes shown in Figure 2 operate in parallel to prevent dataflow bottlenecks, implemented using the Python multiprocessing library. First, volumes enter the acquisition process (Figure 2, left) via scanner connection. The first volume (following dummy scan equilibration) is used as the native EPI for masks. Optionally, *SynthSeg* is applied to the native EPI to identify grey matter (GM), white matter (WM), and cerebrospinal fluid (CSF) masks in the participant whilst they are in the scanner. GM can be isolated downstream for cleaner signal, and WM and CSF are used in a general linear model (GLM) downstream to regress noise from the signal (see Section 2.1.5). As well as *SynthSeg*, a user-defined region of interest (ROI) mask can be registered and transformed from a native space into the native EPI space.

**Figure 2:**
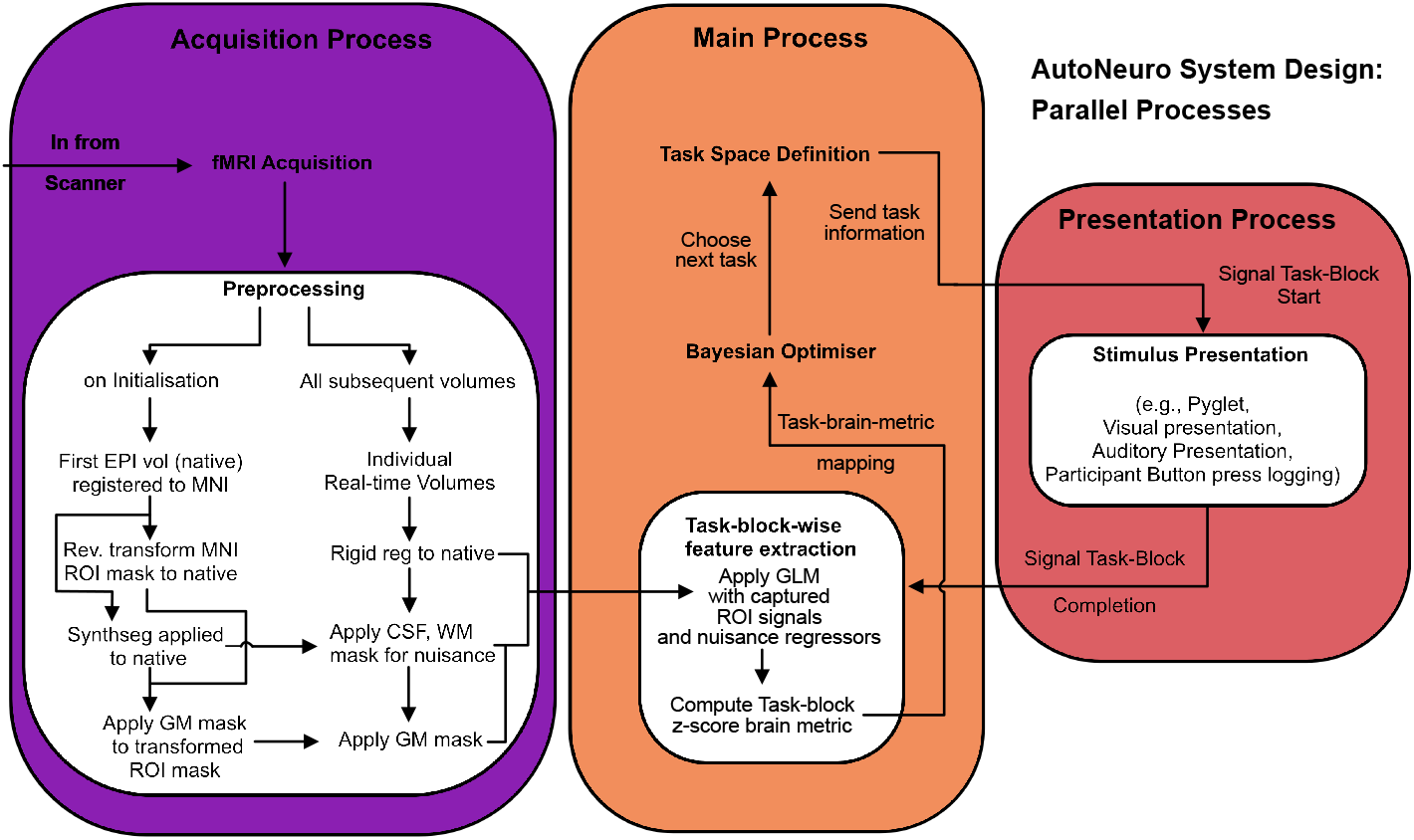
Internal design of AutoNeuro’s workflow. Design is separated by processes that operate in parallel. The three processes are main, acquisition and presentation. The main process dictates task management, the agent (Bayesian optimiser), and timing between processes. The acquisition process receives volumes from the scanner and manages preprocessing before forwarding them to the main process. The presentation process executes task presentation to the participant in the scanner, and communicates block completion to the main process.

Following these steps, the first EPI volume is also used as the registration target for all subsequent volumes. For each incoming volume, the masks described above (which are in EPI native space), can be applied to each incoming volume. This 1., front-loads preprocessing time in the experimental timeline, reducing computational overhead during online operation; and 2., simplifies the registration process to a within-participant rigid transform, rather than the alternative of registering to a standard space using more degrees of freedom.

Each of the now preprocessed volumes is sent to the main process (Figure 2, middle), which counts the number of volumes received for the current experimental block. Upon receiving the expected number of volumes for the current task and/or stimulus, a GLM is applied with nuisance regressors (WM, CSF, motion, drift polynomials, constant; see Section 2.1.5) to compute the user-chosen brain metric (see Section 2.3.4 for demonstration brain metric).

Each experimental block is linked to the brain metric derived from data recorded during that block. The mapping between the observed blocks in the experiment space (see Section 2.1.3) and brain metric is learned by the agent via fitting (Section 2.2). The agent then chooses the next condition in the experimental space to sample based on its learned mapping of the whole space and its internal hyper-parameters. The acquisition function (Section 2.2) is the internal method by which the GP chooses the next task.

Upon choosing the next task, the main process sends task information to the presentation process, beginning the next task block. It also updates the internal state of the acquisition process, which has an internal “block label” state, which associates each incoming volume with the ID of the block that they were captured during. This step acts as an internal clock that keeps volumes and task presentation temporally aligned in the system.

Once a task/stimulus has been chosen, its design is drawn from memory by AutoNeuro and presented to the participant in the presentation process (Figure 2, right). AutoNeuro uses Pyglet, a cross-platform windowing and multimedia library for Python, to present visual stimuli to participants in the presentation process. The toolbox also has the capability of presenting audio stimuli as well as logging button press responses from the participant.

Each possible experimental block is designed and stored in an organised structure of potential experimental conditions before the experiment begins (Section 2.1.3). Since blocks are designed before execution of the task, the duration of the task block is known. The expected number of volumes per experimental block is precomputed, which includes an offsetting of duration to take the haemodynamic response function (HRF) into account.

#### 2.1.3 Experiment Space

To enable the agent to explore the relationship between brain metrics and experimental conditions, we embed all conditions within a shared coordinate space, with axes bounded between limits. We refer to this representation as the “experiment space”. Dimensions of the experiment space can vary by task, by stimulus, or both simultaneously. Each experimental condition is represented by a coordinate within this space, with the number of dimensions in the coordinate system corresponding to the dimensions along which the user wishes to map the experiment space with the chosen brain metric.

A coordinate in the experiment space is therefore an experimental condition that is characterised by where it is situated along each axis of experiment variability. During the experiment, each observed block is temporally bounded within the experiment. In AutoNeuro, the block serves as the sampling unit of the optimisation process: one block yields one summary metric value, which is then treated as a noisy observation of the underlying mapping between experiment space and neural response.

The design of each possible block is defined before data acquisition and is stored in a library for AutoNeuro to pull from in real time. A pre-made design matrix is also associated with each experimental block, which convolves the stimulus presentation window with a double-gamma haemodynamic response function (HRF). The design matrix is used in a subsequent GLM to compute the target brain metric (see Section 2.1.5).

#### 2.1.4 Real-Time Preprocessing

Real-time preprocessing is critical for the AutoNeuro pipeline. Feature extraction and adaptive control must operate on high-quality, spatially aligned data without compromising the temporal constraints of closed-loop experimentation.

For this reason, as much preprocessing as possible is intentionally front-loaded before task presentation begins in order to meet real-time constraints, while preserving signal fidelity.

All points described here demonstrate the current implementation of preprocessing in AutoNeuro, but these steps can be optionally disabled, or replaced with other methods.

During an initialisation phase before task presentation begins, the first acquired EPI volume after dummy volumes are discarded, is registered once to a standard space using the SyNRA (Symmetric Normalisation with Rigid and Affine) algorithm [Avants et al., 2008] from the Advanced Normalisation Tools (ANTs; Tustison et al. [2021]) Python implementation. This step produces a forward transform that maps the participant’s native EPI space to the standard template space. The inverse of this transform can be applied to predefined region of interest (ROI) masks (see Section 2.1.5) to transform them from standard to native space. This ensures that masking of subsequent volumes only requires registration to the native EPI space, which eliminates the computational overhead of warping each new volume to a standard space in real-time.

We then optionally use *SynthSeg* [Billot et al., 2023] to segment the initial EPI volume into grey matter (GM), white matter (WM), and cerebrospinal fluid (CSF). The GM mask is combined with the pre-defined ROIs to isolate grey matter within each. To ensure no overlap between GM, WM, and CSF, both WM and CSF masks are eroded using *scipy*’s binary erosion tool [Virtanen et al., 2020]. The signal from WM and CSF masks is averaged per volume and used to create nuisance regressors for the general linear model (GLM; see Section 2.1.5). Supplementary Figure 1 shows an example participant’s regional GM masks, eroded WM and CSF maps, all outputted from the real-time pipeline.

For each subsequent volume, AutoNeuro first applies a rapid rigid-body registration to the reference EPI volume using ANTs’ *Rigid* transform. This volume is then resampled into alignment with the native EPI using nearest-neighbour interpolation. With subsequent volumes established in the native space, each of the above described masks is applied to each incoming volume.

The ANTs rigid transform for each volume is decomposed into translation and rotation components by reading the forward affine matrix and converting its linear part to Euler angles (xyz order). These six parameters constitute the standard motion regressors, which are used as nuisance regressors in the GLM.

#### 2.1.5 Brain Feature Extraction: Experiment Space Search Metric

This section describes the design and utility of brain metric calculation in the context of the AutoNeuro system. Specifics of the metric used in our demonstration can be found in Section 2.3.4.

Real-time feature extraction is implemented to derive quantitative summaries of ongoing brain activity that could guide adaptive task selection. All metrics are computed online as data accrues following preprocessing (see Section 2.1.4) and are written to a continuously updated history. This history supports both closed-loop decision-making during acquisition and subsequent offline analyses.

At each TR, preprocessed volumes are appended to the existing time series and analysed using a general linear modelling (GLM) approach, applied to the target brain metric. The design matrix incorporates regressors representing the timing of previously presented task blocks together with nuisance covariates to account for structured noise. Model parameters are estimated using ordinary least squares, and statistical contrasts are derived for every completed block. The resulting block-wise estimates are variance-normalised and converted to a common scale so that metrics from different regions and time points are directly comparable.

A GLM is fit independently to the BOLD time series (regional or otherwise) using a design matrix containing task regressors for each presented block and nuisance covariates (2nd order drift polynomial, constant, motion estimate, WM and CSF; see Section 2.1.5). Regression coefficients are estimated using ordinary least squares.

Importantly, the time series of each regressor’s whole time series is updated at the end of each block. The GLM is re-fit repeatedly using every observation in the history of the experiment to improve the signal-to-noise ratio of all observations (including past observations) after each iteration.

### 2.2 Bayesian Optimisation for Adaptive Task Selection

The agent used to guide task selection is within a contained function, meaning prospective users can change the agent to meet various experimental needs. The following is the “default” agent implementation in AutoNeuro; a Bayesian optimiser which utilises a Gaussian Process (GP) regression model.

The objective of the Bayesian optimiser is to minimise the uncertainty around the mapping between experiment space and calculated metric and to efficiently explore the experiment space before exploiting areas of the space that it expects will maximise the target brain metric (see Section 2.3.4).

Following each completed task block, the blocks identifier, timestamp, and the recorded brain metric are recorded and passed to the optimiser as a single block-level observation. At the core of the optimiser is a Gaussian Process (GP) regression model, which provides a probabilistic, non-parametric estimate of the underlying objective function over the experiment space. The GP models the relationship between task parameters *x* and observed performance *f* (*x*), assuming:

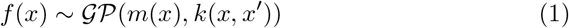

where *m*(*x*) is the mean function (set to zero), and *k*(*x, x*^*′*^) is a covariance (kernel) function that defines the similarity between points in experiment space (for demonstration experiment kernel, see Section 2.3).

The GP is implemented using the *Gaussian Process Regressor* class from *scikit learn* [Pedregosa et al., 2011], with automatic hyperparameter optimisation of the kernel via marginal likelihood maximisation.

The optimiser maintains a continuously updated history of all block-metric pairs (Section 2.3.4). Once a minimum number of samples had been collected, the GP is retrained after each completed task block. Input features are normalised to ensure all task dimensions contribute equally to distance calculations. To improve model robustness, the GP optimisation is repeated multiple times with random restarts. This prevents convergence to local optima in hyperparameter fitting.

### 2.3 Demonstration of Toolbox with Real-Time Data Collection

The goal of this study is to demonstrate the operation, flexibility, and scientific utility of AutoNeuro. We therefore implemented a simple demonstration paradigm designed to show how AutoNeuro can adaptively optimise task parameters to find the optimum of a user-defined neural feature across multiple tasks. Although we describe a specific demonstration implementation, AutoNeuro is designed to enable arbitrary experiment spaces, neural metrics and agent behaviours.

We note that AutoNeuro does not require this specific implementation of slice forwarding, or this specific scanner configuration, and will operate on any data received via the Gadgetron connection.

#### 2.3.1 Experimental Design

For our demonstration, the experiment space comprised five tasks organised along a single abstract dimension. This dimension was constructed by flattening two originally distinct task axes: a motor tapping axis and a visual checkerboard axis. Rather than treating these axes as independent factors, they were combined to form a unified task dimension capturing graded variations in task engagement across modalities.

The five tasks included:

- High opacity checkerboard – a visually stimulating condition with strong stimulus contrast.
- Low opacity checkerboard – a visually less stimulating condition with reduced stimulus contrast.
- Rest block – a baseline condition with no motor or visual demands.
- Slow finger tapping – a motor task performed at a slow pace.
- Fast finger tapping – a motor task performed at a rapid pace.

These tasks were assigned coordinates along the bounded single task dimension to reflect their relative positions in terms of cognitive and sensorimotor demands. Specifically, they were equally spaced along the axis at −1, −0.5, 0, 0.5, and 1. The fast finger tapping task occupied one extreme (*−* 1), representing high motor demand; the high opacity checkerboard occupied the other extreme (*−* 1), representing high visual demand; and the rest block was placed in the middle (0), representing minimal task engagement. The intermediate coordinates corresponded to slow tapping (− 0.5) and low opacity checkerboard (0.5), capturing graded transitions along the task dimension between the extremes of motor and visual engagement. Each task block lasted between 10 and 15 seconds, each being jittered slightly to avoid periodicity.

#### 2.3.2 fMRI Data Acquisition

Four participants (4 male, age range 21-56) were scanned for two 10-minute long sessions each. Eight initial dummy volumes were collected prior to the 10 minute scan to allow signal stabilisation (AutoNeuro receives these dummy volumes and discards them, with the number of volumes ignored determined by the user). Task order was determined online by the optimisation algorithm and therefore was not predetermined by the experimenters. All participants provided written informed consent prior to the experiment, and the study was approved by the local ethics committee at University College London.

Functional images were acquired on a 3T Siemens Prisma scanner using a 32-channel head coil (Birkbeck-UCL Centre for Neuroimaging). Multi-band echo-planar imaging (EPI) was performed with a repetition time (TR) of 1 second, echo time (TE) of 35.2ms, flip angle 60, and 2mm isotropic voxel size. A multi-band factor of 4 was used, with 106 × 106 in-plane matrix and full phase-encoding coverage. Images were reconstructed in real time using the FIRE protocol and forwarded to the processing workstation for online analysis. Head movement was minimised with a head stabilisation device (MR Minimal Motion System, Patent number US2025213412A1; Mangal et al. [2025]). Visual stimuli were presented via a projector onto a rear-projection screen, which participants viewed through an angled mirror mounted on the head coil.

#### 2.3.3 System Acquisition Pipeline

Image data were transferred using the Siemen’s FIRE (Fast Image Reconstruction Environment) protocol, implemented as a generic add-in to the standard CMRR reconstruction sequence. All image reconstruction was performed directly on the scanner. The reconstructed images were outputted in ISMRMRD format and passed to an intermediate PC (configured with Gadgetron; Hansen and Sørensen [2013]) for compatibility, while also being stored in the standard Siemens database.

A lightweight Gadgetron script (see toolbox repository hyperlink here) running on the intermediate PC handled image forwarding, transmitting the reconstructed ISMRMRD-formatted images via TCP/IP to a task-presentation MacMini for subsequent processing. No additional processing was performed at this stage; the transfer script functioned solely to forward reconstructed images in real time. The task-presentation MacMini was used to run AutoNeuro.

AutoNeuro establishes a persistent connection to the listening Gadgetron server via the Gadgetron Python package, receiving reconstructed 3D image slices as numpy arrays as they are produced. The system waits for the expected number of slices, which are appended into a 3D array. When all slices are received for the given TR, the whole volume is processed by downstream steps. The ISMRMRD image format used here includes a header which contains each slices index, ensuring ordering is correct.

#### 2.3.4 Target Brain Metric

Specific to our demonstration experiment, the target brain metric was a regional contrast value that quantified differential task-related activation between two ROIs. The ROIs were derived from activation maps from *Neuroquery* [Dockès et al., 2020], using the labels “checkerboard” and “finger-tapping” (see Figure 4A, top). Each map was thresholded and binarised to create ROIs where the highest activation was located in each map. These masks were applied during preprocessing for each incoming volume (see Section 2.1.4) and voxels within each ROIs grey matter were averaged.

GLM was applied to the ROI time series. Beta coefficients were derived for each task block *i*, regressing out nuisance terms (see Section 2.1.5) for each ROI. The regression coefficient of each ROI for each observed block was converted into a t-statistic using the estimated coefficient variance and residual degrees of freedom. The t-statistic was subsequently transformed into a standard normal z-score using the cumulative distribution function of the Student’s t distribution. The optimisation metric for each block was then defined as the difference between ROI z-scores:

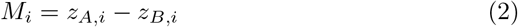

where *M*_*i*_ is the metric for task block *i*, and *A* and *B* indicate each of the two ROIs.

At each update, the metric computation returned the vector of block-wise contrasts 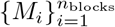 for all blocks presented so far. This formulation yields a variance-normalised, statistically interpretable estimate of differential task engagement between the two regions while accounting for temporal noise structure and nuisance effects. All metric computations were performed online within the temporal resolution of acquisition (see Figure 3).

**Figure 3:**
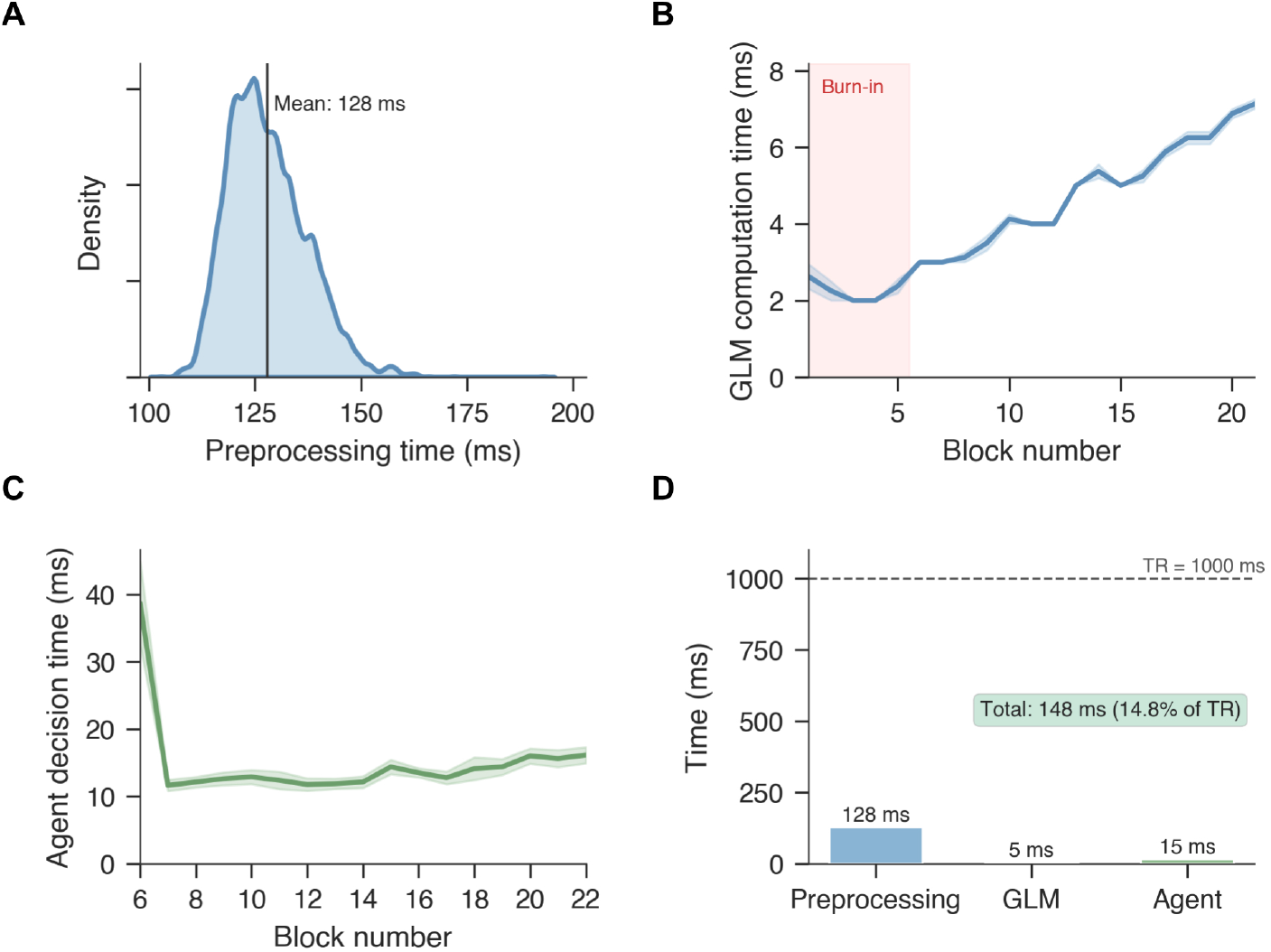
System Performance Benchmark. (A) Density distribution of time taken to preprocess incoming volumes, across runs of all participants. Centre line indicates mean of the distribution. (B) Average duration of GLM computation across runs, per block. Red background indicates the initial “burn-in” period where experimenter-chosen tasks are first presented, before agent decision takes control of task selection. (C) Average duration of agent choosing next task to sample in the task space across task block. Note x-axis begins at block 6 due to the first 5 tasks being chosen by the experimenter (see Section 2.1.3). (D) Total average time to process a volume in the analysis pipeline, separated by stages. Dotted line indicates one TR, at which point volumes would be coming into the pipeline faster than they were getting processed, causing a queue that delays the presentation process.

For each participant, the experiment was conducted twice: once with the target brain metric defined as *M*_*i*_ = *z*_C,*i*_ *− z*_F,*i*_, and once with the sign reversed (*z*_C,*i*_*− z*_F,*i*_), where C and F denote the “checkerboard” and “finger-tapping” masks respectively (Section 2.3.1). This manipulation allowed us to evaluate whether the Bayesian optimiser would systematically shift task selection toward regions of the experiment space that maximise activation in each of the masks during “exploitation”.

#### 2.3.5 Gaussian Process Design

The kernel is the covariance model the GP uses to model the relationship between experiment space and brain metric. A custom sum-kernel was designed as a combination of a White kernel and a Matérn kernel for use in AutoNeuro’s agent (see Section 2.2):

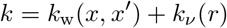

*k*_w_(*x, x*^*′*^) denotes the White kernel, where *x* and *x*^*′*^ denote two coordinates in the experiment space. *k*_*v*_ (*r*) denotes the Matérn kernel.

The White Kernel was initialised with a variance parameter 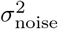 and is used to model uncorrelated noise in observations. Formally, the White Kernel is defined as:

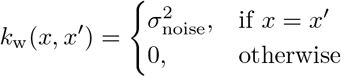

The parameter 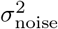 is the signal variance hyperparameter. 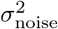is modelled as a bounded hyperparameter, where the noise level is estimated during the fitting process. Here, we initialise 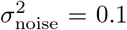, with bounds 1*e −* 8 and 1. The White kernel models independent observation noise by adding variance only when two inputs coincide, ensuring the GP distinguishes structured signal (captured by the Matérn component) from uncorrelated measurement variability.]

For the purposes of the present study, it is sufficient to interpret the Matérn kernel as a distance-based similarity rule with tunable smoothness and correlation range. Specifically, the Matérn kernel is a covariance function that determines how similar neural responses are expected to be for neighbouring points in the experiment space.

Intuitively, the kernel assumes that experimental conditions that are close together in the space (e.g., slow vs. fast tapping) should produce more similar neural responses than task conditions that are far apart (e.g., fast tapping vs. high-contrast checkerboard). The strength and shape of this similarity relationship are governed by two parameters: the length scale *𝓁* and the smoothness parameter *v*. The length scale *𝓁* controls how rapidly similarity decays with distance in the experiment space. A small length scale allows the model to treat nearby task points as relatively independent, producing sharper changes in the predicted response across the experiment space. A large length scale enforces stronger coupling between neighbouring tasks, resulting in smoother variation across the space. The smoothness parameter *v* determines how rough or smooth functions drawn from the Gaussian Process prior are allowed to be. Smaller values of *v* permit rougher, less smooth functions, whereas larger values approach the very smooth behaviour of the squared exponential kernel. We set *v* = 1.5 by default, which corresponds to functions that are once mean-square differentiable. This provides moderate smoothness: the model can interpolate between neighbouring task points while still allowing local irregularities in the neural response landscape.

Formally, the Matérn covariance between two experiment space coordinates *x* and *x* depends only on their Euclidean distance *r* = *xx/𝓁*. The full analytical form of the kernel is given by:

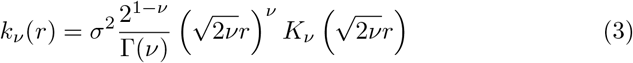

In this expression, *σ*^2^ is the brain metric signal variance and determines the overall amplitude of covariance between experiment space points. Larger values allow greater variation in the latent neural response across the experiment space. The term Γ(*v*) is the Gamma function, which acts as a normalising constant ensuring the kernel remains properly scaled. The function *K*_*v*_ (*·*) denotes the modified Bessel function of the second kind, which governs how covariance decays as the distance *r* increases.

Together, these terms define a covariance that decreases smoothly as the distance between two task coordinates increases. When *r* is small (i.e., nearby points), the covariance is high, implying similar predicted neural responses. As *r* grows, the Bessel function term causes covariance to decay at a rate determined jointly by the length scale *𝓁* and smoothness parameter *v*. For specific values of *v* (including *v* = 1.5), the Matérn kernel simplifies to a closed-form expression, which yields a flexible but not overly smooth function.

#### 2.3.6 Acquisition Function and Task Selection

Coordinate selection in the experiment space is governed by the acquisition function. Here, we utilised the upper confidence bound (UCB) acquisition function, but prospective users of AutoNeuro may alter the chosen acquisition function to suit their needs. The UCB is defined as:

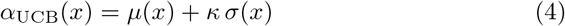

where *µ*(*x*) and *σ*(*x*) denote the GP posterior mean and standard deviation, respectively. The parameter *κ* controls the trade-off between exploration and exploitation by weighting uncertainty relative to expected reward: larger values favour sampling uncertain task points (exploration), whereas smaller values prioritise task points with high predicted mean response (exploitation).

For the present demonstration, we used a heuristic to incrementally decay *κ* as the experiment developed. This is an additional method (alongside the UCB) to shift agent behaviour from explorative (making *σ*(*x*) larger in Equation 4, the models measure of uncertainty) to exploitative (making *µ*(*x*) larger in Equation 4, the models estimate of the landscape). Here, we used a log decay heuristic, defined as:

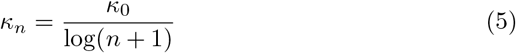

where *κ*_*n*_ denotes the current blocks *κ* value, and *κ*_0_ denotes the *κ* value at the first block.

We initialised *κ*_0_ = 50, a deliberately large value relative to the scale of the posterior mean, ensuring that early task selection is driven primarily by uncertainty rather than predicted response magnitude. This is particularly important in a small, discrete experiment space, where premature exploitation could otherwise lead to early convergence before the GP posterior is sufficiently informed. This approach is particularly useful given the instability of GP model fitting in early blocks, where there is very little data.

The optimisation process runs continuously throughout the experiment without a predefined convergence criterion. At each iteration, the GP posterior is updated with the participant’s latest neural response, along with updated nuisance regressors.

## 3 Results

### 3.1 Real-Time Pipeline Performance and Stability

We first validated the performance and stability of the real-time pipeline. Across runs and participants, the wall-clock processing latency per incoming volume remained well below the 1*s* TR.

There were two computationally-unique conditions under which performance was assessed. First, is preprocessing that was executed once per incoming volume (see Section 2.1.4). These types of computation required on average 128ms (Figure 3A), with no single volume exceeding 190ms across all runs and participants.

Second, are block-level computations; timepoints at which summary measures of the block were computed (Section 2.1.5); and Gaussian process (GP) training and task selection computations (Section 2.3.5). At these block boundaries, the system transitions into a brief “idle” block that serves as a synchronisation interval, ensuring that all computations complete before the onset of the next task block. Across participants, both GLM estimation and GP-based task selection required no more than 20ms at block completion (Figure 3B, C). From the first block onward, GLM computation time increased modestly as the number of regressors in the design matrix grew with each completed block (Figure 3B), with average duration rising from approximately 2.5ms to approximately 7ms across 20 blocks. The initial GP model training (at block 6, after the 5 initial burn-in blocks; Figure 3C) required longer (mean 40ms), whereas subsequent updates required less than 20ms per block.

Combining preprocessing (128ms on average) with block-boundary computations (20ms), the maximum observed processing load per TR was approximately 148ms, corresponding to 14.8% of the 1s TR. At no point did processing approach the TR duration, preventing volume queuing and ensuring uninterrupted closed-loop task adaptation. These results indicate substantial headroom for incorporating more computationally intensive analysis procedures in future extensions of AutoNeuro.

### 3.2 Real-Time Preprocessing

Figure 4 shows the real-time preprocessing and analysis (see Section 2.1.5) that is applied at TR resolution in an example participant run. Figure 4A shows the native space EPI of the individual, with the top panel showing where the grey matter segmentation overlapped with the ROI masks. The colours of each of the ROIs correspond to the values in Figure 4B, which show how the summary z-score output of the general linear model applied at each experimental block. The CSF and white matter masks shown in the middle and bottom panels of Figure 4A are example masks that were averaged across and used as nuisance regressors in the GLM. Colours in these masks correspond to those in Figure 4C, where an example time series of each is shown.

**Figure 4:**
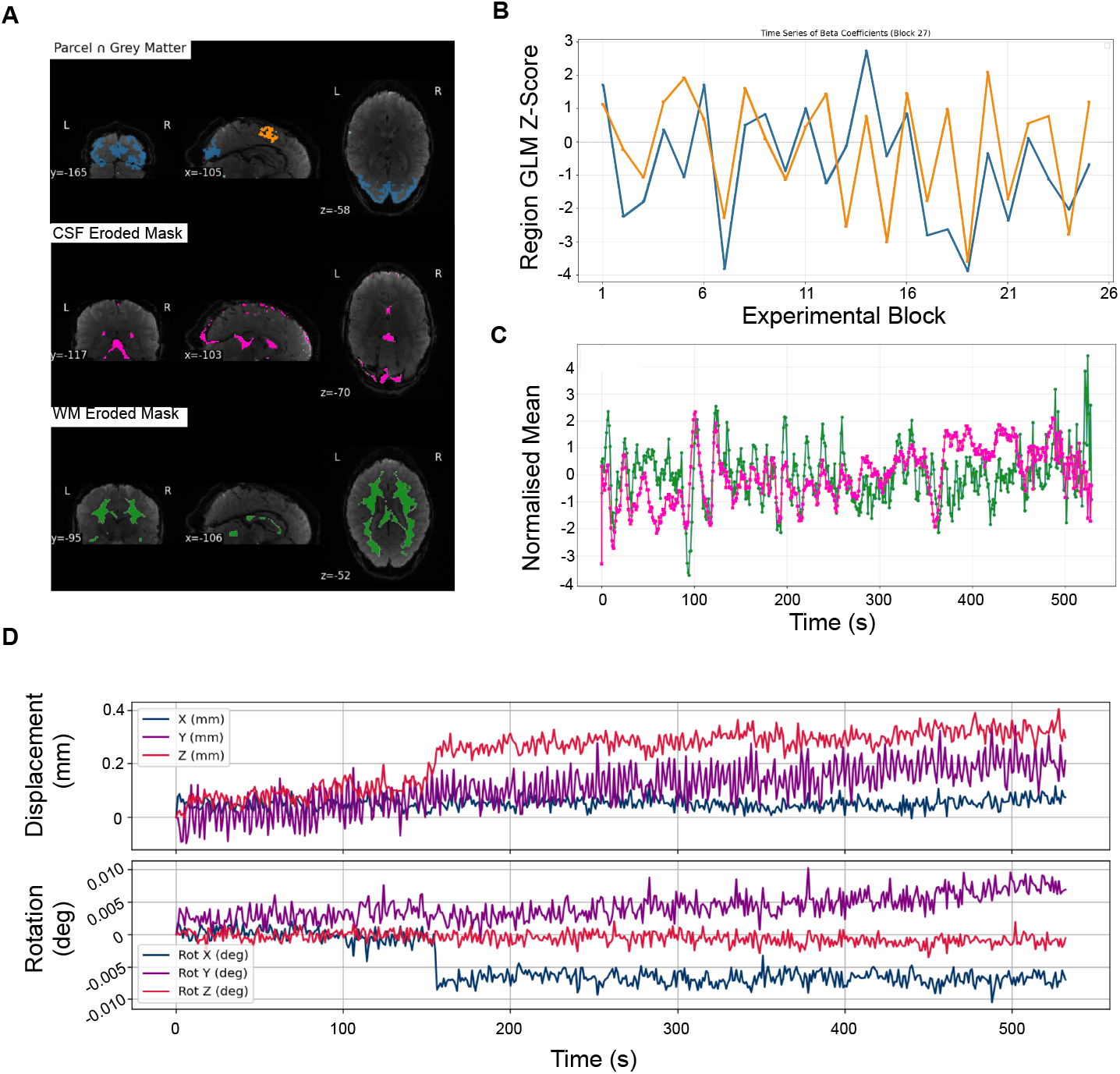
Example Participant Online Preprocessing and Analysis. **A** Example participant native volume with each mask applied. The top panel shows the overlap of the grey matter mask and the motor and visual masks, coloured orange and blue respectively. The eroded CSF mask is shown in pink, and the eroded white matter mask in green. **B** The output z-score of each region after 26 task blocks. A single z-value is computed for each region, for each experimental block following fitting of the GLM. Colours correspond to the masks in A. **C** Normalised mean values within the CSF and white matter masks, each used as nuisance regressors in the GLM. Colours correspond to masks in **D** Estimated absolute translational and rotational motion from the example participant, calculated from registration process. Each of the six measures is used in the GLM as a nuisance regressor.

Figure 4D then shows the motion estimate computed from registration to the first native volume. All preprocessing shown here was computed in real-time during the experiment, with the difference in values between regions in Figure 4B being the brain metric that was mapped onto the experimental space (see Equation 2).

### 3.3 Region-Based Contrast Demonstration

With the real-time pipeline validated, we next examined how AutoNeuro adapted task selection based on the relationship between the target brain metric and the experiment space. Each participant initially completed a “burn-in” period of five experimenter-chosen tasks, during which all five task conditions were presented once (Figure 5A, highlighted). Following this, the Bayesian optimiser began selecting tasks adaptively, informed by prior observations of the brain metric and the Gaussian Process model.

**Figure 5:**
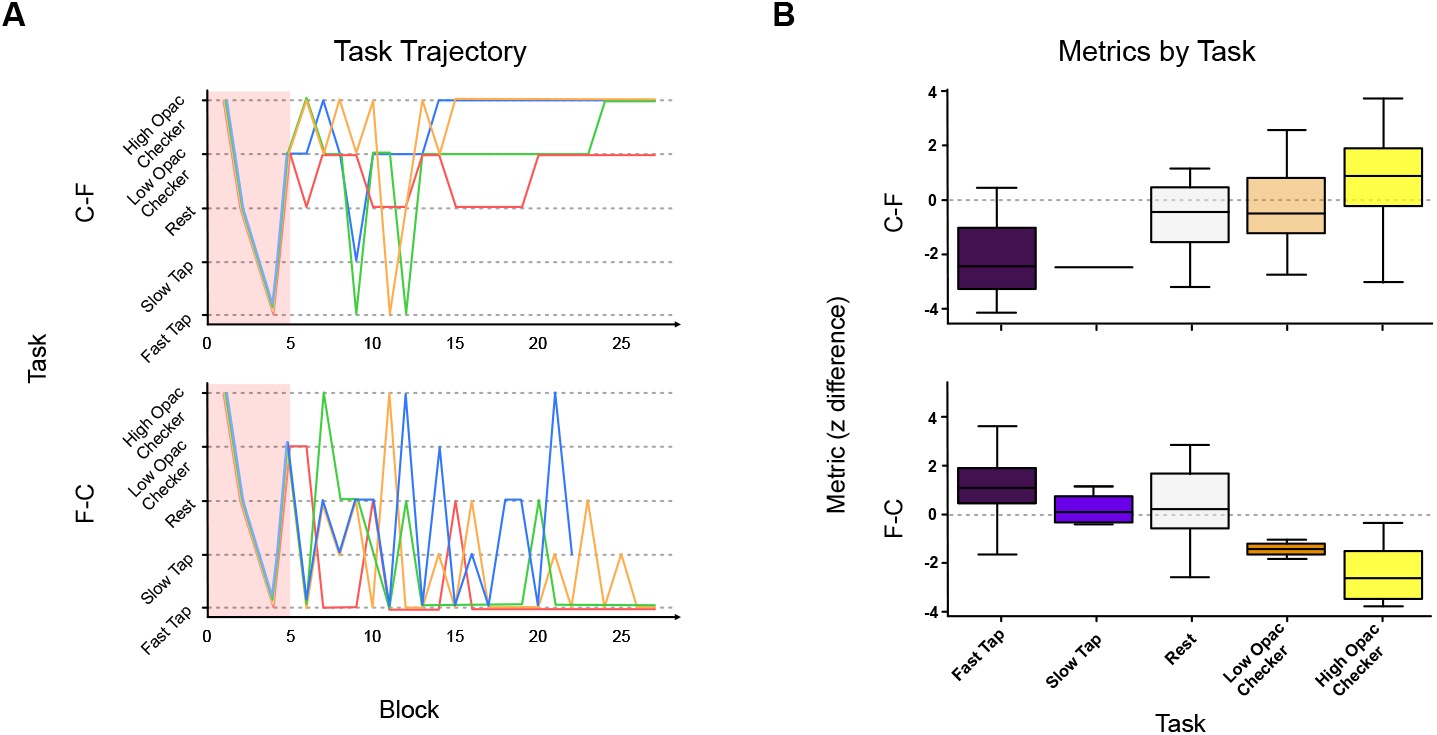
Adaptive Maximisation of Regional Contrast. (**A**) Task trajectories over time for all participants. Each line represents a participant, with colour differentiating subjects. The top panel uses the metric *z*_C_ *− z*_F_, the bottom panel uses *z*_F_*− z*_C_. The first five blocks (highlighted) constitute the experimenter-chosen burn-in period, after which AutoNeuro’s Bayesian optimiser adaptively selected tasks based on the observed brain metric. Trajectories show convergence toward task conditions that maximise the respective target metric. (**B**) Block-wise measured metric values corresponding to each task type across participants. More blue indicates motor heavy task, more red indicates more visually intense task.

Each participant performed two runs, corresponding to the two metrics. These are: *z*_C_*− z*_F_ and *z*_F_*− z*_C_, which determine the two differences in activation between the two regions (see Figure 4 top, and Section 2.1.5).

In the top panel of Figure 5A, we show the trajectory of tasks selected when maximising *z*_C_ *− z*_F_, and in the bottom panel, the trajectory for maximising *z*_F_*− z*_C_. Across participants, we observe that the optimiser transitions from initial exploration to exploitation, converging on task conditions that maximise the target metric. Specifically, maximising *z*_C_ *− z*_F_ favoured visual checker-board tasks over motor tapping tasks, whereas maximising *z*_F_*− z*_C_ favoured the opposite pattern.

Figure 5B presents the measured block-wise metrics across the experiment, across participants. This illustrates the relationship between task space and observed metric values that AutoNeuro leverages during adaptive task selection. Together, these results demonstrate that the framework successfully searches the task space exploratively (initially varying trajectories in Figure 5A) before identifying and converging on task conditions that elicit maximal regional contrast, confirming the neuroadaptive capabilities of the system.

### 3.4 Gaussian Processes Modelling of the Experiment Space

To examine how the GP model captured the relationship between experiment space and the measured brain metric, we analysed the model posterior, specifically the predicted mean and standard deviation at each task point. Figure 6A presents the projected mapping for each participant and each brain metric condition for each run. The final GP model exhibited a clear capability to map the contrast of motor and visual tasks onto visual and motor regional differences.

**Figure 6:**
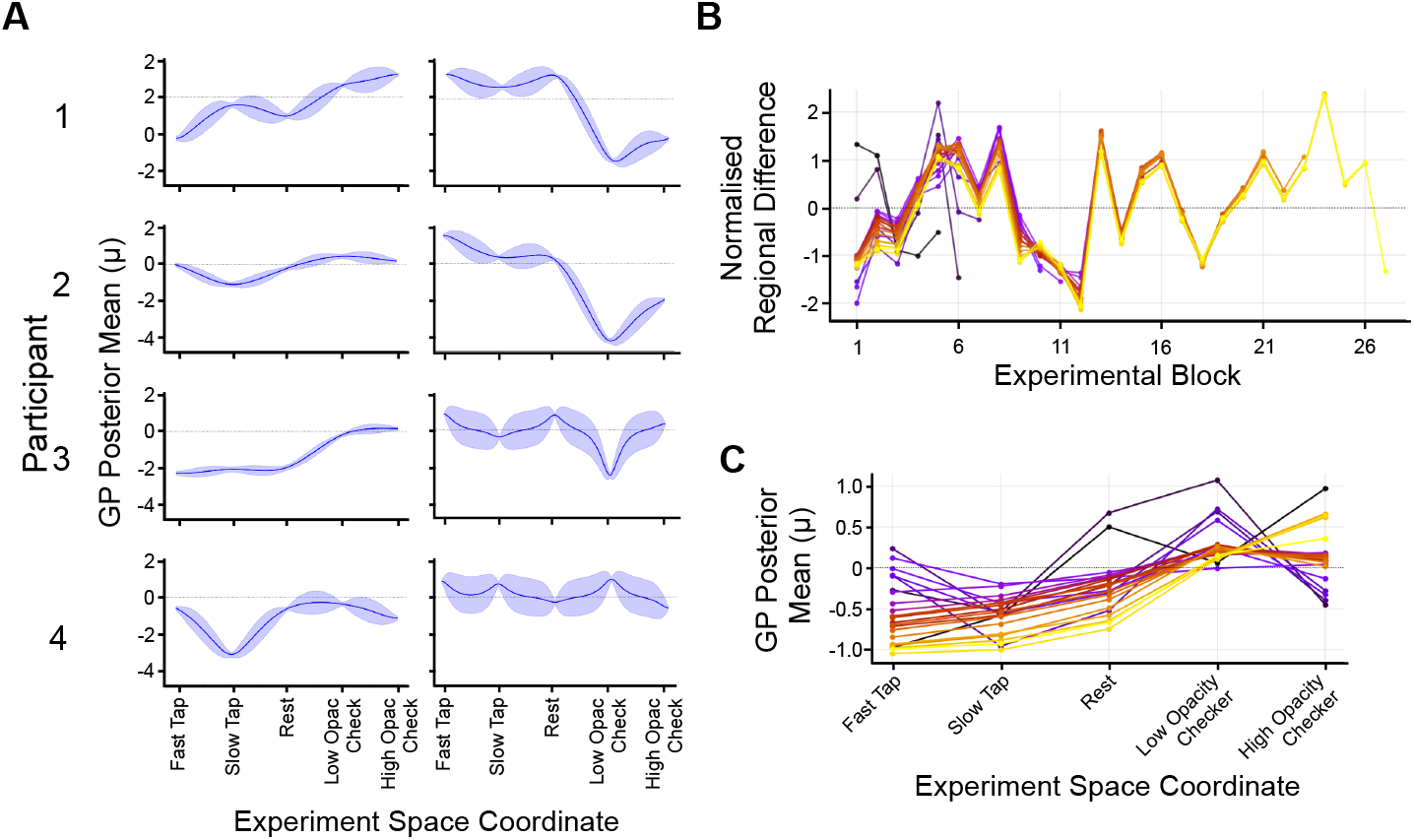
GP Posterior. (A) The end GP posterior of each participant, in each condition. Line shows the posterior mean, shaded area indicates the models’ uncertainty of the space. Rows show each participant, columns show run’s target brain metric: region difference of C-F on the left, and F-C on the right. X-axis shows the experiment space. (B) Participant 3’s C-F run’s regional difference value. Colour of line indicates the experiment block, with yellow being later in a run. Note how early observations change over time as more data is acquired. (C) Development of GP posterior for Participant 3’s C-F run as blocks developed. Colour of line indicates the experiment block, with yellow being later in a run.

Figures 6B and C show an example run of participant 3, C-F, that demonstrates how the GP learns the mapping over time. At the end of each block during the experiment, both the GLM (Figure 6B) and GP model (Figure 6C) are re-fitted to the data, meaning not only is the most recently completed task block appended to the fitted data, but the history of observations is also updated in the context of the whole experiment observed so far.

## 4 Discussion

We present AutoNeuro, a practical, robust toolbox for neuroadaptive experimental design in human neuroimaging, and demonstrate its use in a proof-of-concept study.

Here, we demonstrated that AutoNeuro operates within the temporal constraints of online acquisition, completing image capture, preprocessing, feature estimation, model inference, and task selection within a 1s TR (Figure 3). The framework successfully leveraged a brain-derived metric to first explore a user-defined experiment space and then steer the experiment toward conditions that maximised that metric (Figure 5), illustrating the feasibility of closed-loop neuroadaptive experimentation.

Several existing tools support components of real-time fMRI workflows (see Supplementary Table 1). OpenNFT provides an open-source platform for neurofeedback [Koush et al., 2017], while the AFNI real-time plug-in enables online acquisition, motion correction, and ROI time-course extraction [Cox, 1996]. Turbo-BrainVoyager offers comparable functionality within a commercial environment [Goebel, 2012]. AutoNeuro differs in purpose: rather than delivering feedback or monitoring signals, it embeds adaptive experimental control directly into the acquisition loop, allowing neural responses to determine which task is presented next. This shifts real-time fMRI from pre-specified paradigms toward data-driven exploration of brain–task relationships.

The integration of Bayesian optimisation provides a principled mechanism for this exploration. By balancing sampling of uncertain task points in the model with exploitation of promising conditions, the framework is well suited to noisy neural data. Importantly, Bayesian optimisation yields a surrogate model of the response surface, enabling characterisation of how neural activity varies across the task space rather than merely identifying a single optimum.

A constraint in the default implementation is that the experiment space must be discretised into temporally finite task blocks. Although the underlying cognitive or stimulus dimensions may be continuous, the optimiser requires discrete observations to update the Gaussian Process posterior. This discretisation defines the granularity at which the neural objective can be estimated and directly influences optimisation performance. Notably, block duration introduces a fundamental trade-off between statistical reliability and search efficiency. Longer blocks yield more volumes per condition, increasing the precision of the GLM estimate and improving the signal-to-noise ratio of the derived brain metric. This reduces observation noise in the GP model and can stabilise hyperparameter estimation. However, longer blocks reduce the number of conditions sampled within a fixed scanning time, limiting exploration of the experiment space and slowing convergence. Conversely, shorter blocks permit denser sampling and more rapid posterior updating, but at the cost of noisier metric estimates, potentially benefitting from the interpolation inherent in GP models. Future work is needed to explore these trade-offs systematically.

Our initial demonstration was modest in scope, involving four participants and a simplified experiment space and metric. The intention was not to map a novel brain–task relationship, but to illustrate the neuroadaptive process. The true benefits of neuroadaptive approach is clearer over much larger experimental spaces that are uncertain and hard to sample exhaustively. The framework is flexible: the brain metric need not be a regional contrast, and could instead capture connectivity, entropy, or behavioural measures for example. Similarly, different optimisation strategies (e.g., multi-armed bandits) may be more suitable for larger or more complex discrete experiment spaces. Alternative deep learning frameworks may also be useful to replace the GP or Bayesian Optimiser in some situations, e.g., Neural Processes [Garnelo et al., 2018], PFNs4BO [Müller et al., 2023] or GPTOpt [Meindl et al., 2026].

Additionally, the acquisition function utilised in the model need not be an Upper Confidence Bound—which emphasises the explore–exploit behaviour demonstrated here—but could instead aim to maximise variance in sampling or identify tasks that reduce entropy.

During our implementation of the toolbox, we found that the choice of brain metric and task space critically shapes the experiment. Metrics define the “lens” through which neural responses are interpreted, and convergence in metric space may not fully reflect underlying cognitive or clinical phenomena. This highlights the potential for multi-finality: the same brain-metric value can result from multiple, distinct task configurations, so convergence in metric space may not uniquely reflect the underlying cognitive or neural processes. Similarly, defining experiment space along predetermined axes assumes that relevant variation occurs within those dimensions, potentially overlooking unmodelled processes. While these constraints limit post-hoc exploration, they also enforce principled experimental design, preventing biased analyses where hypotheses are formulated after observing the data [Lorenz et al., 2021].

Future work will focus on extending AutoNeuro’s capabilities. Incorporating multi-objective optimisation would allow simultaneous optimisation of multiple outcome measures, such as neural activity and behavioural performance, which is particularly relevant for clinical or translational applications [Yang et al., 2019]. This demonstration of AutoNeuro’s capabilities in a low-dimensional task space provides a foundation for future work applying similar approaches to larger task spaces. Such approaches can enable efficient, participant-relevant exploration of task spaces that would be too large to search using a traditional grid search. The open-source, modular design provides a foundation for implementing new metrics, optimisation strategies, and experimental paradigms, supporting a broad range of neuroadaptive research beyond conventional static designs.

In summary, AutoNeuro demonstrates the feasibility of real-time neuroadaptive experiments. By enabling neural responses to guide task selection, it supports data-driven exploration of brain–task relationships while maintaining a flexible, robust framework that can be adapted for diverse applications.

## 5 Data and Code Availability

The AutoNeuro Python framework is an open-access tool that can be found here. The wiki contains instructions for set-up both offline and online to connect the system to a scanner.

The collected data and output results for this proof-of-concept study can be found on OpenNeuro, here.

## Ethics Statement

All participants provided written informed consent prior to the experiment, and the study was approved by the local ethics committee at University College London.

## Author Contributions

**FD**: Conceptualisation, Methodology, Investigation, Resources, Visualisation, Writing – Review and Editing, Supervision, Project Administration, Funding Acquisition. **DH**: Methodology, Software, Validation, Formal Analysis, Investigation, Resources, Data Curation, Writing – Original Draft, Writing – Review and Analysis, Visualisation, Project Administration. **RL**: Conceptualisation, Methodology, Software, Validation, Formal Analysis, Investigation, Supervision. **RR**: Investigation, Project Administration. **OS**: Methodology, Software, Validation, Formal Analysis, Data Curation, Writing – Original Draft, Writing – Review and Analysis, Visualisation.

