## Supplemental Information for "AutoNeuro: An Open-Source fMRI Toolbox for Real-Time Neuroadaptive Task Design"

### Supplementary Information

| Framework | License | Language | Maintenance | Imaging Integration | Preprocessing Quality | Feature Scope | Task / Feedback Control | Adaptive Logic | Ease of Use |
|---|---|---|---|---|---|---|---|---|---|
| Pyneal [1] | Open-source | Python | Active | fMRI DICOM stream (Siemens/GE/Philips) | Motion, de-trending | ROI / MVPA (plugin-based) | Local socket / GUI | – | Simple setup, clear docs |
| OpenNFT / pyOpenNFT [2] | Open-source | MATLAB, Python | Active | fMRI DICOM / TCP | Full SPM-based pipeline | ROI / Connectivity / MVPA | GUI + Psychtoolbox link | – | MATLAB API |
| RT-Cloud [3] | Open-source | Python | Active | fMRI DICOM → Cloud API (BIDS-RT) | User-defined via custom scripts | Flexible (MVPA, GLM, etc.) | HTTP or socket messaging | – | Requires Docker/cloud setup |
| RTPSpy [4] | Open-source | Python | Active | fMRI Local DICOM stream | Advanced (motion, smoothing, regressors) | ROI / custom features | GUI + simulation mode | – | – |
| AFNI Real-Time [5] | Open-source | C, TCL | Active | fMRI DICOM / socket connection | Motion + GLM in real-time | ROI / GLM only | User scripting | – | Clear docs, existing frontends |
| FRIEND Engine [6] | Open-source | C++, Python | Limited | fMRI DICOM / FSL backend | Yes (motion, filtering) | ROI / Connectivity / MVPA | GUI | Neurofeedback focused | Windows build |
| Turbo-BrainVoyager [7] | Commercial | C++ | Active | Vendor-integrated | Full pipeline | ROI / GLM / MVPA | GUI + BV Studio | – | Closed ecosystem |
| improv [8] | Open-source | Python | Limited | 2P Calcium Imaging | Full Pipeline | Python API | Bayesian optimiser | Turn-key | |
| AutoNeuro | Open-source | Python | Active | Gadgetron, TCP/IP, ICE | Modular low-latency chain | Arbitrary, user-defined | Python API | Bayesian optimiser | Turn-key, simulator |

Table 1: Comparison of real-time fMRI frameworks.

[7] Brain Innovation - Products - Turbo-BrainVoyager. https://www.brainvoyager.com/products//products/turbobrainvoyager.html.

[8] Anne Draelos, Matthew D. Loring, Maxim Nikitchenko, Chaichontat Sriworarat, Pranjal Gupta, Daniel Y. Sprague, Eftychios Pnevmatikakis, Andrea Giovannucci, Tyler Benster, Karl Deisseroth, John M. Pearson, and Eva A. Naumann. A software platform for real-time and adaptive neuroscience experiments. *Nat Commun*, 16(1):9909, November 2025.
